# Detecting remote homolog using structure alignment algorithms and machine learning

**DOI:** 10.1101/2022.12.15.520536

**Authors:** Priscila Caroline de Sousa Costa, Tetsu Sakamoto

## Abstract

Remote homolog detection is a classic problem in Bioinformatics. It attempts to identify distantly related proteins sharing a similar structure. Methods that can accurately detect remote homologs benefit protein functional annotation. Recent computational advances in methods predicting the three-dimensional structure of a protein from amino acid sequences allow the massive use of structural data to develop new tools for identifying remote homologs. In this work, we created a discriminative SVM-based method based on structural alignment algorithms (FATCAT, TM-Align, and LovoAlign) to detect whether a protein is a remote homolog with any proteins in the SCOPe database. The final model showed a ROC AUC of 0.9191.

## Introduction

Proteins have diverse functionalities and are fundamental in several, if not all, biological processes. Thus, functionally annotating each of them helps us understand the processes in which they are involved. Common procedures to carry out the functional annotation of proteins are based on sequence alignment methods, in which a sequence similarity search is performed in databases to look for functionally characterized proteins with high sequence similarity with the queried proteins. Although efficient and widely used, when two functionally similar proteins share an identity below 40% (twilight zone) these methods are barely able to detect their evolutionary or functional relationships (Chung and Subbiah, 1996; Laurents *et al*., 1994). Since protein function is defined primarily by its structure, it is common to find proteins sharing high structure similarity despite low sequence identity. For those protein pairs, we call Remote Homologous.

One widely used database to assist the development of methods for identifying remote homologs is SCOP (Hubbard *et al*., 1997) and SCOPe (Fox *et al*., 2014). In these databases, proteins with known structures are organized in a hierarchical structure according to their evolutionary relationship. Proteins that belong to the same SCOP family have clear evidence about their evolutionary origin. Proteins from one SCOP superfamily but from different families are more distantly related, and the similarity between them is only detectable at the structural level. Within the context of the SCOP classification, proteins from the same superfamily but from a different family are considered remote homologs.

Correctly identifying remote homologs is of great relevance for the protein functional annotation (Lobb *et al*., 2015), and several tools have been developed to perform this task (Wan and Xu, 2005; Fariselli *et al*., 2007; Chen *et al*., 2018). Most of them seek to extract features that allow the discrimination of remote homologs from the amino acid sequence, as it is the most abundant data type (Steinegger *et al*., 2019; Li *et al*., 2017). It is known that remote homolog identification using structural data results in better performance (Wan and Xu, 2005). However, the restricted number of structural data available, due to the methodological limitation in obtaining experimental data on protein structures, has restricted until now the development and application of tools that use this kind of data. This scenario, fortunately, tends to change with the current advance in computational methods that accurately predict the three-dimensional structure of proteins from amino acid sequences (Jumper *et al*., 2021; Lin *et al*., 2022). These advances promise to reduce the large discrepancy between the volume of sequence data and structural data. Therefore, it is interesting to rethink the development of such software that takes advantage of the structure data.

In this work, we ensembled some structural alignment algorithms and used their results to compose the dataset to train a classification model that identifies whether a protein is a remote homolog with a protein in the SCOPe database. To the best of our knowledge, we consider this work to be one of the first to explore the use of structural alignment tools for the identification of remote homologs.

## Material and Methods

### Data source

We used SCOPe (v.2.08) (Fox *et al*., 2014) as the data source. In this database, we obtained the three-dimensional structural data of the proteins in PDB format and information about the superfamilies to which each protein is classified. In this work, we have used the subset of the proteins that share less than 95% identity between them according to the ASTRAL compendium (Fox *et al*., 2014). This subset will be referred to as SCOPe 95 and consists of 35,494 proteins classified into 2,065 superfamilies.

### Building the dataset

To create the dataset used to build the classifier, firstly we randomly selected a representative protein for each superfamily found in SCOPe 95 (2,065 superfamilies). Then, each representative protein was structurally aligned with all proteins in SCOPe 95 (subject) using the software FATCAT (Li *et al*., 2020). The metrics that describe the structure alignment (RMSD, score, alignment size) were used as variables for classification (Table 1). The alignment results were ranked according to the score provided by the software. Then, each pair of query and subject proteins were identified whether they are remote homologs (class 1) or not (class 0) by checking whether they belong to the same SCOPe superfamily.

**Table 1:** Descriptive statistics of the dataset for each class

| features | Remote homologs (N=8,457) |  |  |  |  |  |  | Non remote homologs (N=20,650) |  |  |  |  |  |  |
| --- | --- | --- | --- | --- | --- | --- | --- | --- | --- | --- | --- | --- | --- | --- |
|  | mean | std | min | 25% | 50% | 75% | max | mean | std | min | 25% | 50% | 75% | max |
| lovo_finalScore | 0.62 | 0.21 | 0.02 | 0.46 | 0.64 | 0.80 | 1.00 | 0.25 | 0.11 | 0.01 | 0.18 | 0.24 | 0.30 | 0.86 |
| lovo_coverage | 139.38 | 99.42 | 14.00 | 76.00 | 111.00 | 174.00 | 1299.00 | 108.02 | 68.64 | 12.00 | 64.00 | 90.00 | 129.00 | 781.00 |
| lovo_rmsd | 4.00 | 3.74 | 0.10 | 1.83 | 2.87 | 4.67 | 42.47 | 9.38 | 5.09 | 0.60 | 5.77 | 8.53 | 11.90 | 61.58 |
| lovo_gaps | 10.66 | 9.25 | 0.00 | 4.00 | 9.00 | 15.00 | 104.00 | 17.47 | 9.79 | 0.00 | 11.00 | 16.00 | 22.00 | 85.00 |
| lovo_finalScoreNorm | 0.73 | 0.17 | 0.19 | 0.61 | 0.75 | 0.87 | 1.00 | 0.48 | 0.15 | 0.14 | 0.36 | 0.45 | 0.58 | 0.99 |
| tm_AliLen | 134.53 | 97.09 | 10.00 | 72.00 | 108.00 | 167.00 | 1246.00 | 82.40 | 48.80 | 5.00 | 51.00 | 71.00 | 98.00 | 452.00 |
| tm_RMSD | 2.47 | 1.08 | 0.10 | 1.69 | 2.38 | 3.14 | 10.03 | 4.35 | 1.22 | 0.33 | 3.54 | 4.35 | 5.12 | 9.70 |
| tm_n_ident/n_aln | 0.29 | 0.23 | 0.00 | 0.12 | 0.22 | 0.39 | 1.00 | 0.07 | 0.04 | 0.00 | 0.05 | 0.07 | 0.09 | 0.52 |
| tm_TM-score (chain 2) | 0.68 | 0.19 | 0.05 | 0.54 | 0.69 | 0.83 | 1.00 | 0.27 | 0.13 | 0.02 | 0.19 | 0.26 | 0.33 | 0.92 |
| tm_d0 (chain 2) | 4.43 | 1.54 | 0.50 | 3.38 | 4.26 | 5.40 | 12.41 | 5.35 | 1.74 | 0.50 | 4.02 | 5.32 | 6.49 | 12.41 |
| fatcat_subject-len | 166.75 | 118.61 | 17.00 | 89.00 | 132.00 | 212.00 | 1521.00 | 242.45 | 169.19 | 20.00 | 119.00 | 205.00 | 314.00 | 1521.00 |
| fatcat_Twists | 0.45 | 1.02 | 0.00 | 0.00 | 0.00 | 0.00 | 5.00 | 1.72 | 1.63 | 0.00 | 0.00 | 1.00 | 3.00 | 5.00 |
| fatcat_ini-len | 117.03 | 90.25 | 8.00 | 64.00 | 88.00 | 144.00 | 1104.00 | 74.80 | 61.97 | 8.00 | 32.00 | 56.00 | 96.00 | 616.00 |
| fatcat_ini-rmsd | 2.42 | 1.50 | 0.08 | 1.45 | 2.18 | 3.13 | 28.20 | 3.57 | 2.15 | 0.08 | 2.31 | 3.14 | 4.15 | 30.22 |
| fatcat_opt-equ | 134.12 | 96.81 | 8.00 | 72.00 | 107.00 | 166.00 | 1147.00 | 85.26 | 64.69 | 2.00 | 42.00 | 69.00 | 107.00 | 692.00 |
| fatcat_opt-rmsd | 2.38 | 0.89 | 0.10 | 1.70 | 2.57 | 3.06 | 11.40 | 3.06 | 1.00 | 0.02 | 2.64 | 3.06 | 3.34 | 16.07 |
| fatcat_chain-rmsd | 3.70 | 4.04 | 0.08 | 1.46 | 2.26 | 3.89 | 40.82 | 10.06 | 7.13 | 0.08 | 3.65 | 9.33 | 14.70 | 48.20 |
| fatcat_Score | 270.54 | 241.44 | 17.56 | 118.53 | 196.92 | 333.46 | 2410.83 | 124.72 | 97.41 | 11.06 | 56.74 | 100.01 | 157.98 | 829.12 |
| fatcat_align-len | 166.39 | 125.12 | 8.00 | 85.00 | 131.00 | 214.00 | 1832.00 | 143.44 | 119.22 | 2.00 | 67.00 | 111.00 | 181.00 | 1381.00 |
| fatcat_Gaps | 32.27 | 50.12 | 0.00 | 4.00 | 15.00 | 42.00 | 1006.00 | 58.18 | 62.00 | 0.00 | 17.00 | 41.00 | 78.00 | 853.00 |
| fatcat_rel_score | 0.58 | 0.25 | 0.02 | 0.37 | 0.58 | 0.79 | 1.00 | 0.21 | 0.13 | 0.01 | 0.11 | 0.17 | 0.27 | 1.00 |
| fatcat_rel_align | 0.82 | 0.18 | 0.06 | 0.73 | 0.88 | 0.96 | 1.00 | 0.40 | 0.21 | 0.01 | 0.23 | 0.38 | 0.54 | 1.00 |

Since the data is highly unbalanced, because more protein pairs are not remote homologs compared to those that are, we performed a filter step, in which we selected the alignments of a couple of protein pairs to build the classifier. In this procedure, among the alignment results obtained for a protein representing a superfamily, we selected up to five alignments between remote homologs that had the worst scores, and up to five alignments between non-remote homologs that had the best scores. Additionally, alignment results of at least 10 protein pairs (five remote homologous pairs and five remote non-homologous pairs) were randomly chosen and included in the dataset. In the end, the final dataset comprised the alignment results of 29,108 protein pairs.

Subsequently, these 29,108 protein pairs were submitted to two more structural alignment programs: TM-align (Zhang and Skolnick, 2005)and LovoAlign (Martínez *et al*., 2007). The metrics provided by these aligners were also added to the dataset and used as variables. After removing the missing data rows, the final dataset contained 29,107 rows and 22 columns (Table 1).

### Building the classifier

Machine learning methods were applied using Python language (v.3.8) and Scikit-learn library (v.1.2.0) (Pedregosa *et al*., 2011). The metric used to evaluate the performance of each generated model was the area under the ROC curve (ROC AUC) since it is robust even when dealing with unbalanced data. In addition, we submitted the standardized data before submitting the data for training. In the first instance, we applied different supervised machine learning methods (SVM, decision tree, and logistic regression) to the dataset and evaluated it with cross-validation (k=10). The best-performing method was selected for the hyperparameter tuning step. For the fitting, approximately 10% of the data set was separated to be used as validation (validation set). The rest of the dataset was submitted to the tuning, which was conducted using the grid search approach with cross-validation (k=10). Among the models evaluated during the tuning, the 10 best-performing models were selected and tested with the validation set. The model that performed the best in this last test was chosen as the final model. It is important to highlight that in all steps involving the partitioning of the data set (cross-validation and separation of the validation set), the partitioning occurred in such a way that the results of alignments involving the same query protein are always in the same partition. This procedure helps verify the generalization of the model in classifying superfamilies that are not present in the training set.

## Results

### Building classifier for remote homolog detection

To find a suitable method to model the remote homolog classification problem proposed in this work, we applied our dataset to different supervised machine learning methods. Among the methods applied are SVM, Decision Trees, and Logistic Regression. To compare the performance of each model, cross-validation (k=10) and the ROC AUC metric were used. In addition, each of the variables present in the dataset was standardized, and all models were trained using the default value of their hyperparameters. Among the methods tested, the model trained with SVM showed the best performance (ROC AUC: 0.9048) and was then chosen to perform the hyperparameter tuning.

For hyperparameter tuning, approximately 10% of the data were separated for validation and the rest were used for the fitting. The fitting followed the grid search approach with cross-validation (k=10). The hyperparameters of the SVM method and their respective values tested were: C (1, 10, 100, 1000), gamma (0.1, 0.01, 0.001, 0.0001), kernel (rbf, linear, poly, sigmoid), and class_weigth (balanced). In total, 84 models were evaluated. The 10 models that had the highest ROC AUC values when tested with the validation set, had ROC AUC values between 0.9203 and 0.9317. The model that had the highest ROC AUC value with the validation data was chosen to be the final model and had the following hyperparameter values: kernel: rbf; C: 1000; gamma: 0.001; class_weight: “balanced”. Other model performance metrics were obtained after performing cross-validation (k=10) on the entire dataset (Table 2). The final model presented a ROC AUC of 0.9191.

**Table 2:** Performance metrics of the final model

| **Metric** | **Mean** | **Standard deviation** |
| --- | --- | --- |
| precision | 0.8015 | 0.0199 |
| recall | 0.9331 | 0.0125 |
| f1-score | 0.8622 | 0.0149 |
| accuracy | 0.9133 | 0.01 |
| rocauc | 0.9191 | 0.0099 |

### Model performance for each SCOPe superfamily

The final model was then evaluated for each of the 2,065 superfamilies represented in the dataset. This was performed using the leave-one-group-out approach of cross-validation, where one group contains only the alignment results of one representative of a SCOPe superfamily as a query. For each group evaluated, the accuracy was calculated. In this analysis, we could verify that out of the 2,065 superfamilies evaluated, 1,603 superfamilies (77.62%) showed an accuracy above 90%, and 1,910 superfamilies (92.5%) had an accuracy above 70% (Figure 1). Most of the superfamilies with an accuracy below 70% are of class “All beta proteins” (with 61 superfamilies), followed by “Alpha and beta proteins (a+b)” (with 33 superfamilies), “Alpha and beta proteins (a/b)” (with 27 superfamilies) and the “All alpha proteins” (with 22 superfamilies).

**Figure 1:**
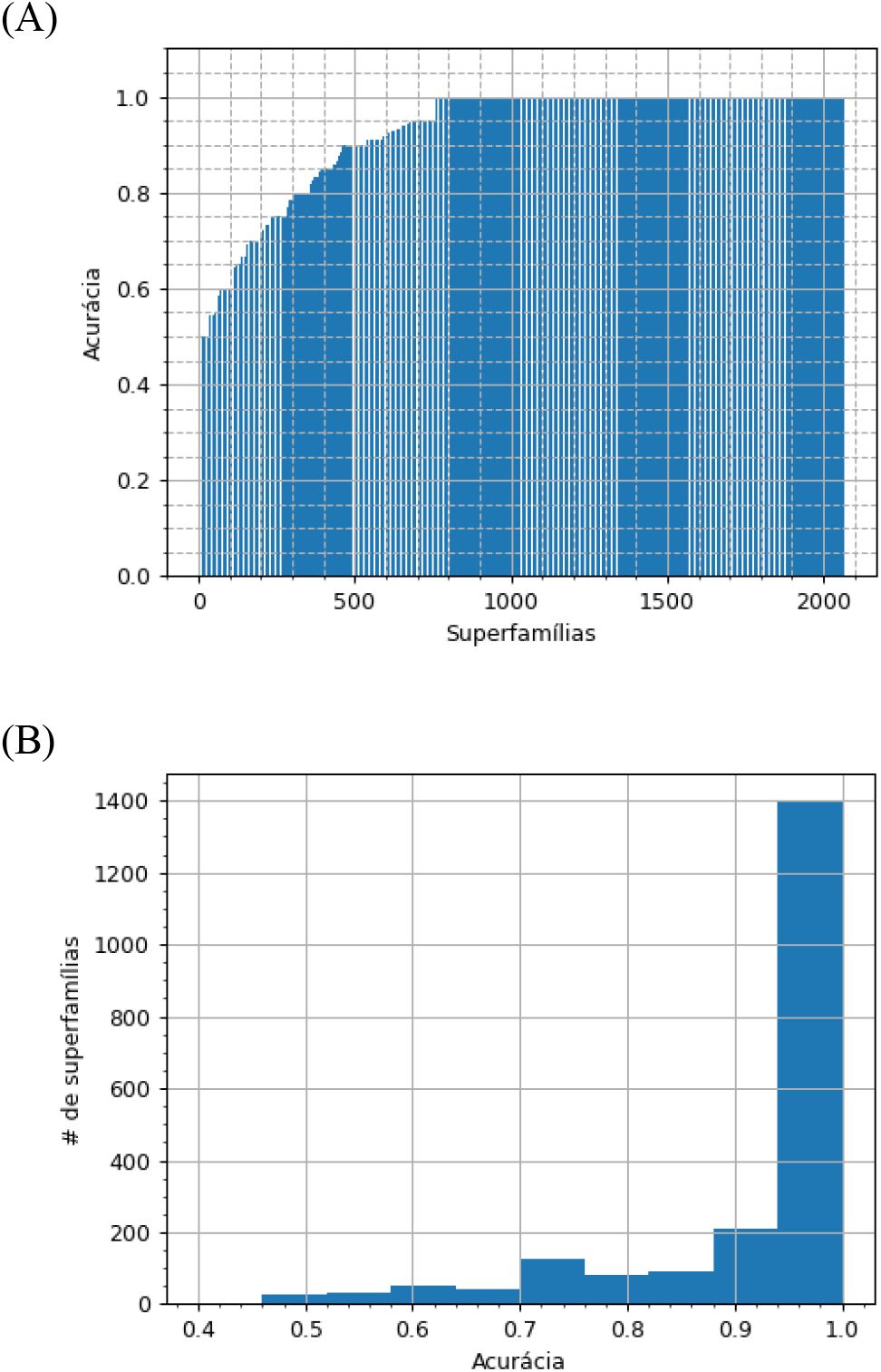
(A) Accuracy and (B) histogram of the accuracy of the final model for each group representative of 2,065 SCOPe superfamilies.

## Discussion

The model created in this work can be categorized as a discriminative method since we treat the problem with discrimination algorithms. However, the approach used in this work differs from most of the methods included in this category. One such difference is that the discriminative methods seek to generate a classifier for each (super)family. In this work, only a single classifier was generated. Our classifier verifies whether a query protein could be a remote homologous with a protein present in the SCOPe database. Another difference is the dataset used to train the model. While discriminative models in general are trained only on (super)families that have representativeness (Chen *et al*., 2018), the approach used in this work allows our model to be trained with almost all alignment results of a pair of proteins. Discriminative but generalized approaches like this work can be found in the Homolog Induction (Karwath and King, 2002) and SVM-HUSTTLE (Shah *et al*., 2008) software.

A widely used dataset to compare discriminative methods is the SCOP-D (Liu *et al*., 2014, 2008). This dataset consists of proteins from the SCOPe v.1.59 database that share an e-value less than 1e-25. In this dataset, there are 54 families and 23 superfamilies that have enough members to generate a classifier. For this dataset, we can find methods that reach a ROC AUC of 0.982, which is the case of the SW-PSSM software (Rangwala and Karypis, 2005). Differences in the approach conducted in this work prevent direct comparison of the performance of our model with other discriminative methods. The model developed in this work reached a lower ROC AUC (0.9191) at the first glance, but it offers the advantage of covering all SCOPe superfamilies. It is noteworthy that the criteria used to build the dataset also offer a more challenging scenario for the classifier. Even with this result, the model reached higher performance than several other models described in the literature (Chen *et al*., 2018).

Despite the general high performance achieved by the model, there are some superfamilies (7.5%) whose model showed a low predictive capacity. For these superfamilies, there is a need for further study to verify the cause of this low performance. One possibility is that the model is dealing with very similar superfamilies, which may be the case for superfamilies of the “All beta proteins” class. In these cases, a solution is to verify the possibility of merging some of these superfamilies or even generating a different classifier for this class of proteins. Another possibility would be to elaborate or extract new variables that could help distinguish these superfamilies. We can also explore new variables using other structural alignment programs, such as DALI (Holm and Sander, 1995), Fast (Zhu and Weng, 2005), and Mammoth (Ortiz *et al*., 2002).

